# Characterization of age-associated inflammasome activation reveals tissue specific differences in transcriptional and post-translational inflammatory responses

**DOI:** 10.1101/2024.04.22.589283

**Authors:** Sarah Talley, Tyler Nguyen, Lily Van Ye, Rasa Valiauga, Jake DeCarlo, Jabra Mustafa, Benjamin Cook, Fletcher A White, Edward M. Campbell

## Abstract

Aging is associated with systemic chronic, low-grade inflammation, termed ‘inflammaging’. This pattern of inflammation is multifactorial and is driven by numerous inflammatory pathways, including the inflammasome. However, most studies to date have examined changes in the transcriptomes that are associated with aging and inflammaging, despite the fact that inflammasome activation is driven by a series of post-translational activation steps, culminating in the cleavage and activation of caspase-1. Here, we utilized transgenic mice expressing a caspase-1 biosensor to examine age-associated inflammasome activation in various organs and tissues to define these post-translational manifestations of inflammaging. Consistent with other studies, we observe increased inflammation, including inflammasome activation, in tissues. However, we note that the degree of inflammasome activation is not uniformly correlated with transcriptional changes commonly used as a surrogate for inflammasome activation in tissues. Furthermore, we used a skull thinning technique to monitor central nervous system inflammasome activation *in vivo* in aged mice and found that neuroinflammation is significantly amplified in aged mice in response to endotoxin challenge. Together, these data reveal that inflammaging is associated with both transcriptional and post-translational inflammatory pathways that are not uniform between tissues and establish new methodologies for measuring age-associated inflammasome activation *in vivo* and *ex vivo*.

## Introduction

Biological aging is associated with a series of age-associated pathologies that reduce organismal fitness and increase susceptibility to numerous diseases that are more prevalent in aging populations. The hallmarks of aging include genomic instability, altered intercellular communication, cellular senescence, mitochondrial dysfunction, deregulated nutrient-sensing and metabolic derangements (1, 2). Because of these attributes, aging has been identified as a risk factor for numerous diseases (3). Some common age-associated diseases include degenerative diseases of the nervous system, cardiovascular disease, cancer, immune system disease, and musculoskeletal disorders (4-14). Aging increases the susceptibility of individuals to these different diseases and infections due to “immunosenescence,” an age-related deterioration of both the innate and adaptive immune system (15). The rise in age-associated diseases is becoming a burden on public health programs, especially in developed countries, necessitating further research on how to combat the development of such diseases (16).

Mechanistically, aging leads to the accumulation of different tissue-specific age-associated phenotypes, accompanied by a state of chronic, low-grade inflammation called “inflammaging” (17). These changes are known to be driven by alterations in the transcriptomes of aging cells, including upregulation of innate immune receptors and genes associated with interferon signaling (18, 19). In many cases, expression of such genes leads to the constitutive release of inflammatory cytokines which contribute to the pathology of inflammaging. This pathology is often associated with the presence of damage associated molecular patterns (DAMPs), which include macromolecules and cellular debris released in response to oxidative stress, other forms of cellular dysfunction and cell death. Sensing of DAMPs by pattern recognition receptors (PRRs) can induce the activation of multi-protein complexes called inflammasomes (20, 21) and constitute post-translational cellular response pathways whereby inactive precursor proteins are cleaved, activated and collectively drive the release of active, inflammatory mediators. Numerous PRRs, including NOD-like receptors (NLRs) and AIM2-like receptors (ALRs), form inflammasome complexes in response to DAMP recognition, cellular injury and stress (22, 23). Following assembly of these multi-protein complexes, inflammasome receptor oligomerization leads to the autoproteolytic cleavage and activation of caspase-1 (22). Activated caspase-1 cleaves and thereby activates inflammatory cytokines, such as pro-IL-1β and pro-IL-18 which lead to propagation of the cellular inflammatory responses (24). Caspase-1 also cleaves gasdermin D (GSDMD) into its functional form, promoting its membrane association and pore formation, which facilitate the release of inflammatory cytokines and can also promote inflammatory cell death known as pyroptosis (24-27).

The most widely studied inflammasome is NLRP3. The activation of the NLRP3 inflammasome canonically involves a two-signal mechanism. An initial priming step (signal 1), occurs in response to PRR recognition of pathogen associated molecular patterns (PAMPs) or cytokine signaling, leading to NF-κB-dependent upregulation of various inflammatory transcripts, including NLRP3 and pro-IL-1β (28). Priming of the cell enhances the responsiveness to inflammasome activating stimuli (signal 2). NLRP3 activation can be induced by lysosomal damage, mitochondrial dysfunction, reactive oxygen species and dysregulation of cytoplasmic ion concentrations, including increases in Ca^2+^ or K^+^ efflux (29-33).

Genetic studies reveal that deletion of *NLRP3* ameliorates some aspects of age-related functional decline (1, 34-36). However, for technical reasons, most studies of inflammaging have largely focused on transcriptional changes of *NLRP3* and other genes that participate in inflammatory responses. Yet, given that inflammasome activation is mediated through a series of post-translational protease cleavage events of caspase-1 and other proteins, monitoring inflammasome activation, rather than changes in inflammasome associated transcripts, is necessary to fully understand the inflammatory sequelae associated with inflammaging.

To characterize global and tissue-specific changes in inflammasome activation during the process of aging, we utilized a transgenic mouse model in which a luciferase-based caspase-1 biosensor is constitively expressed in all tissues (37). Proteolytic activation of caspase-1 thus induces luciferase activation in specific tissues, allowing inflammasome activation to be monitored *in vivo* and *ex vivo* (37-43). These studies reveal a tissue-specific degree of overlap or discordance between age-associated inflammasome activation and upregulation of inflammatory transcripts commonly associated with inflammaging. We observe that inflammasome activation is not directly correlated with transcriptional upregulation of innate immune receptors or inflammatory cytokines. Additionally, using a model of endotoxemia, we establish a system to monitor age-associated changes in inflammasome activation in the central nervous system (CNS), which allows for more effective monitoring of post-translational inflammatory responses in the CNS.

## Results

### Aged Mice Exhibit Tissue-Specific Increases in Inflammasome Activation and Immunoglobulin Deposition

To characterize global changes in inflammasome activation during aging, we used caspase-1 reporter mice described previously to visualize caspase-1 activation *in vivo* and *ex vivo* (37, 40, 41, 43). At ∼18 months of age, mice exhibited increased biosensor activation *in vivo* (**Fig 1A-B**), consistent with age-dependent increase in low-grade systemic inflammation. To identify the source of this increased biosensor signal, we extracted tissues from young (6–12-week-old) and aged (18–24-month-old) mice and biosensor activation was assessed *ex vivo*. We found elevated biosensor activation in some tissues from aged mice, including the pancreas, kidneys, heart, and brain, while biosensor signal remained unchanged in other tissues, such as the lungs and intestines (**Fig 1C, 1D**).

**Fig. 1.**
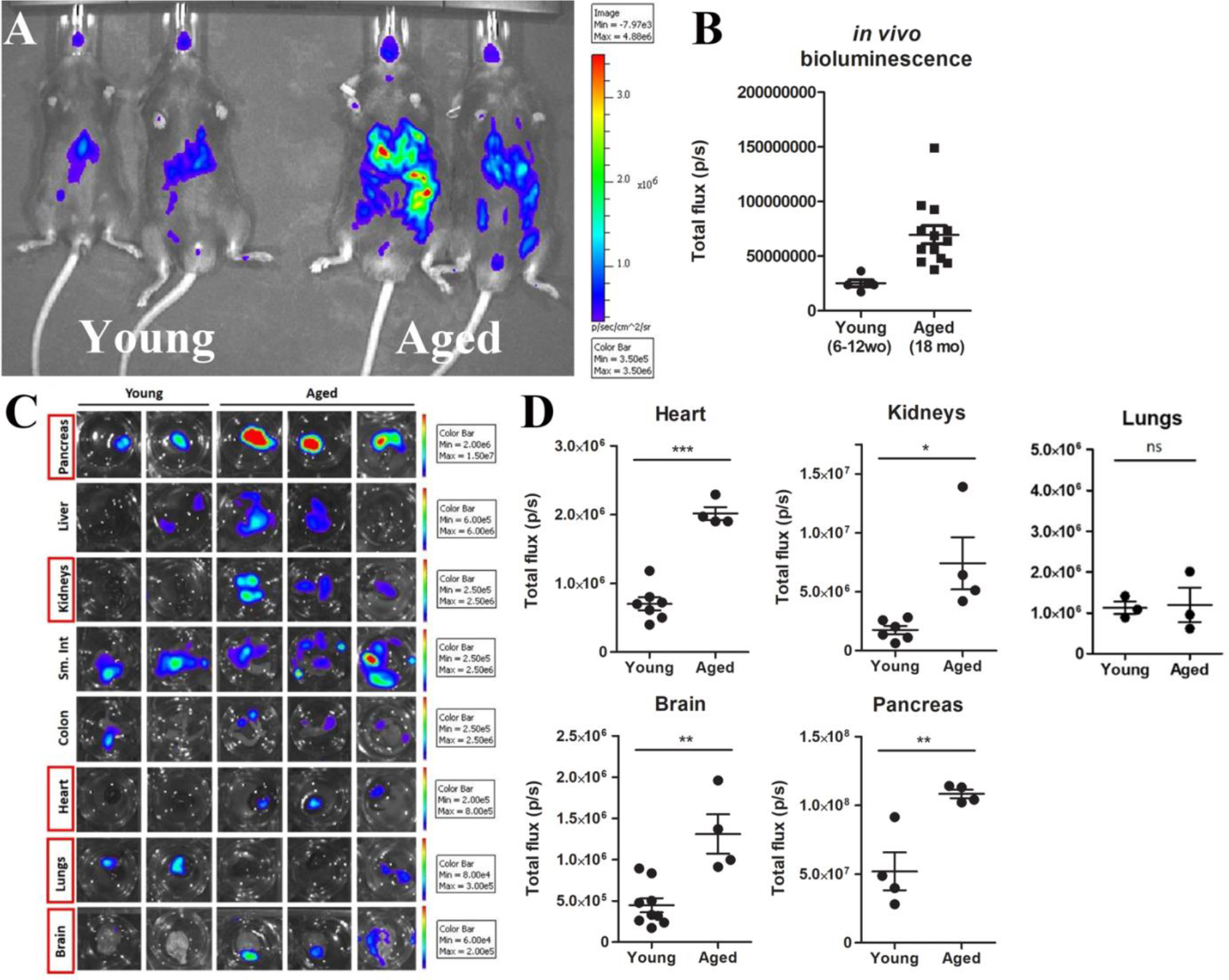
Caspase-1 activation increases *in vivo* during aging and in some tissues *ex vivo*. Representative images and quantification of *in vivo* caspase-1 biosensor activation in young (12 week old) and aging (18-24 month old) mice (A-B). Tissues were extracted from young and aged mice and tissue bioluminescence was measured *ex vivo* (C-D). * = p<0.05, ** = p<0.005, *** = p<0.0005 Student’s *t* test.

To confirm caspase-1 activation in these tissues, we attempted to corroborate biosensor signal by monitoring caspase-1 cleavage via western blot. However, the presence of murine immunoglobulins, particularly in aged tissue, complicated this assessment. Specifically, we used a commonly utilized mouse antibody to caspase-1 that can detect both the uncleaved p45 and cleaved p20 band of caspase-1. Consistent with our biosensor measurements, we observed apparent increases in the amount of cleaved caspase-1 in the brain, heart and kidneys (**Fig S1)**. However, we also observed an increase in cleaved caspase-1 in the lungs, where we measured relatively little age-associated caspase-1 activation using our biosensor (**Fig S1A)**. However, when lysates were analyzed using only the goat α-mouse secondary antibody, we noted bands of similar size and intensity to what was seen when the primary antibody was included, suggesting that much if not all of the observed signal by western blot was due to the presence of murine immunoglobulins in these tissues (**Fig S1B**). We therefore attempted to immunodeplete immunoglobulins from these tissues with protein A/G beads. However, immunodepletion did not effectively remove the 20 kDa band from these lysates (**Fig S2**), which prevented us from assessing the amount of cleaved caspase-1 in aged tissues. Assessment of age-associated immunoglobulins in various organs revealed that this phenomenon was not exclusive to IgG, as both IgM and IgA levels were also increased in many organs (**Fig S3**). Less deposition of all three immunoglobulins was observed in the brain, although an age-dependent deposition of IgG and IgM was still clearly observed (**Fig S3**).

As the NLRP3 inflammasome has been previously implicated in age-associated inflammation (1, 34-36), we treated aged mice with the NLRP3 inhibitor MCC950 to determine the degree to which increased biosensor activation observed in aged mice was derived from increased NLRP3 inflammasome activation (44). Treatment of aged mice with MCC950 induced a significant reduction in biosensor signal emanating from the head, abdomen and paws, reducing total biosensor signal by 50% or greater in these tissues (**Fig 2A, 2B**). A similar trend was noted in the chest, but this difference was not statistically significant (**Fig 2B**).

**Fig. 2:**
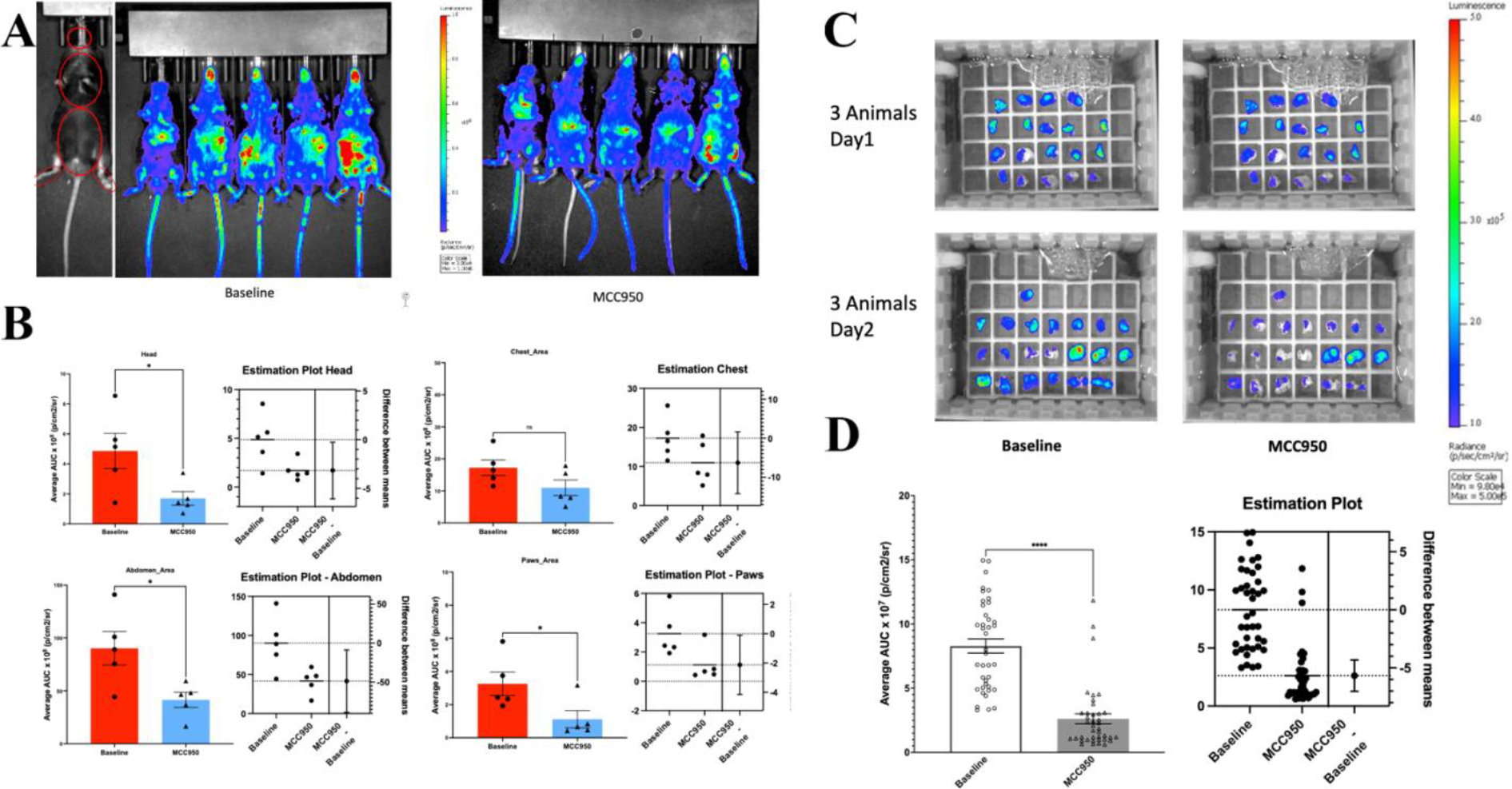
MCC950 attenuated caspase-1 mediated inflammatory signal in aging mice. (A) Sample images show aged caspase-1 activation reporter mouse (22-24 months old) bioluminescence signals before and after MCC950 treatment. MCC950 was treated within 24 hours prior to the second imaging session. The analyzed ROIs (red circles) were quantified across both time points. (B, C) Significantly elevated head, abdominal, and paw caspase-1 activation associated signals in aged mice and significantly decrease post MCC950 treatment. n = 5 mice per group; *p<0.05 paired T-test). (C) CNS brain slices

We similarly monitored biosensor activation in the brain *ex vivo* using a slice culture model. We imaged brain slice cultures from 6 aged animals, applied MCC950 to these cultures and measured biosensor activation again 24 hours after addition of MCC950 (**Fig 2C**). Similar to what we observed following MCC950 administration *in vivo*, we noted a significant decrease in biosensor activation in brain slice cultures following addition of MCC950 (**Fig 2C, 2D**). Collectively, these data demonstrate that the increased biosensor activation observed in aged mice is due to NLRP3 inflammasome activation.

### Aging associated changes in inflammatory transcripts are distinct from changes in inflammasome activation

Next, we assessed changes in commonly measured inflammatory transcripts in tissues of young and aged mice. Notably, although we observed a statistically significant increase of caspase-1 activity in the brain, kidneys, and heart, these changes did not correlate with similar differences in inflammasome related transcripts. Specifically, we did not observe statistically significant changes in *NLRP3* or *IL-18* in these tissues, although a trend towards an increase in *NLRP3* was observed in the kidneys and heart, as well as in the lungs, despite the lungs not exhibiting increased caspase-1 biosensor activation in aged animals (**Fig 3**). We observed statistically significant increases in *pro-IL-1β* and *caspase-1* transcripts in some organs, with the largest and most significant increases observed in the lungs (**Fig 3**). Smaller but significant increases in *pro-IL-1β* and *caspase-1* were observed in the heart and a statistically significant increase in *caspase-1* transcripts was observed in the brain. No significant increase in *pro-IL-1β* transcripts was observed in the brain and no inflammasome associated transcripts were increased in the kidney, despite the increase in *caspase-1* biosensor observed in aged kidneys. Analysis of other inflammatory genes, including *IL-6* and interferon stimulated genes, revealed relatively modest changes in transcription in most organs, with significant increases in interferon stimulated genes *ISG15* and *IFI16* noted in the brain and *ISG15* in the kidneys. Together, these data demonstrate that tissue-specific changes in inflammasome-specific transcripts do not correlate with the increases in caspase-1 activation in aged mice.

**Fig. 3.**
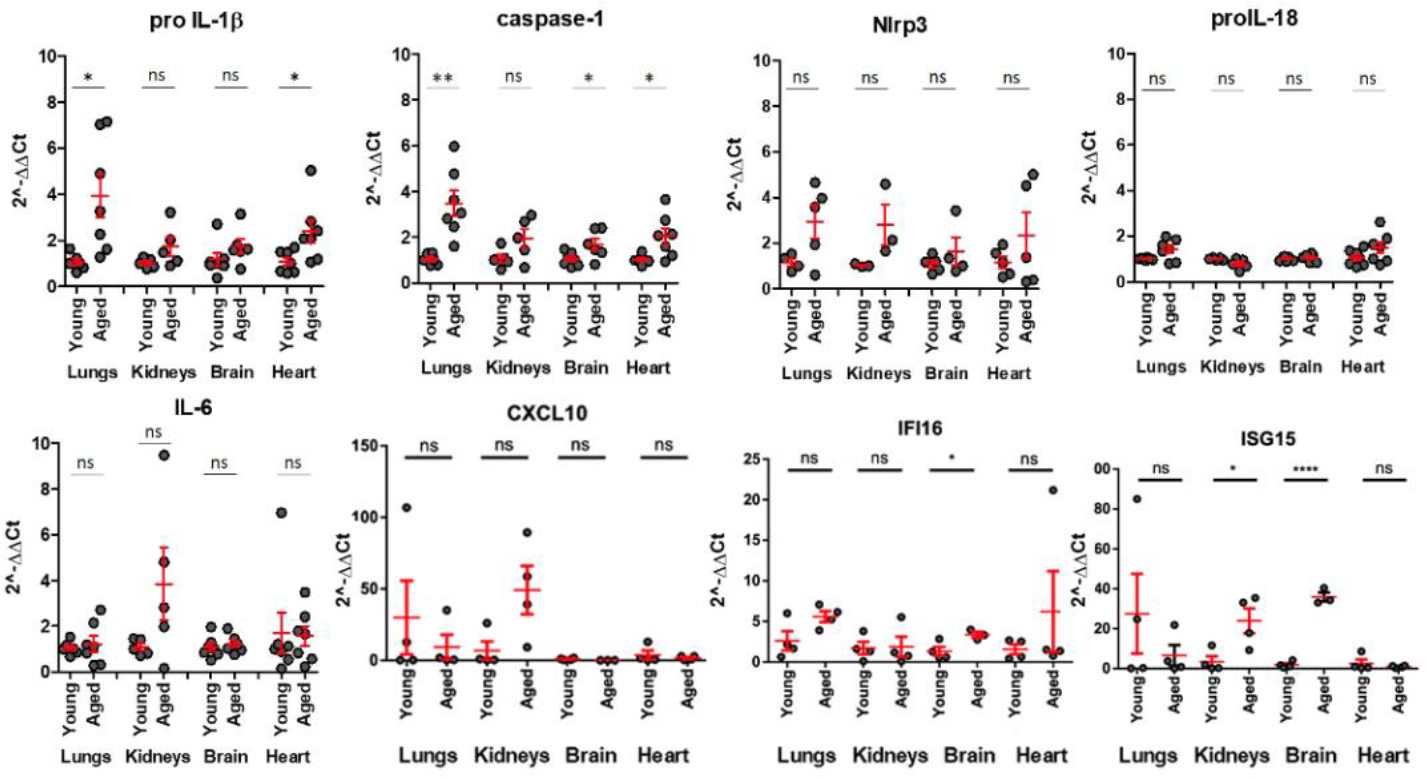
Aged tissues exhibit minimal changes in proinflammatory transcript expression. RNA was isolated from lungs, kidneys, heart, and brain tissues from young (6-12 week old) and aged (18-24 month old) mice and proinflammatory transcript expression was quantified by qRT-PCR. * = p<0.05, ** = p<0.005, **** = p<0.0001 Student’s *t* test.

### Aging aggravates endotoxemia-induced inflammasome activation in the CNS

We next asked if age influenced inflammasome activation in response to endotoxemia, a common model of sepsis. It is known that peripheral administration of lipopolysaccharide (LPS) induces systemic and neuroinflammation through activation of immune cells in the periphery and the release of cytokines and chemokines (45, 46). Age is known to increase the inflammatory and cognitive sequelae induced during sepsis in both humans and mouse models (47, 48). However, most studies of CNS responses to LPS injection have been unable to directly monitor inflammasome activation, instead relying on changes in transcription that may or may not be indicative of inflammasome activation in the CNS. We therefore injected young and aged mice with LPS, and caspase-1 activation and transcriptional changes were measured in tissues 24h later. We detected a robust increase in biosensor activation and upregulation of inflammatory transcripts, most notably *IL-6*, in all tissues (**Fig S4**). We observed a significant increase in biosensor activation following LPS injection in both young and aged mice, specifically in the kidney, brain, heart and lungs, despite a relatively small number of aged biosensor mice included in this experiment (n=3) (Fig 4).

**Fig. 4:**
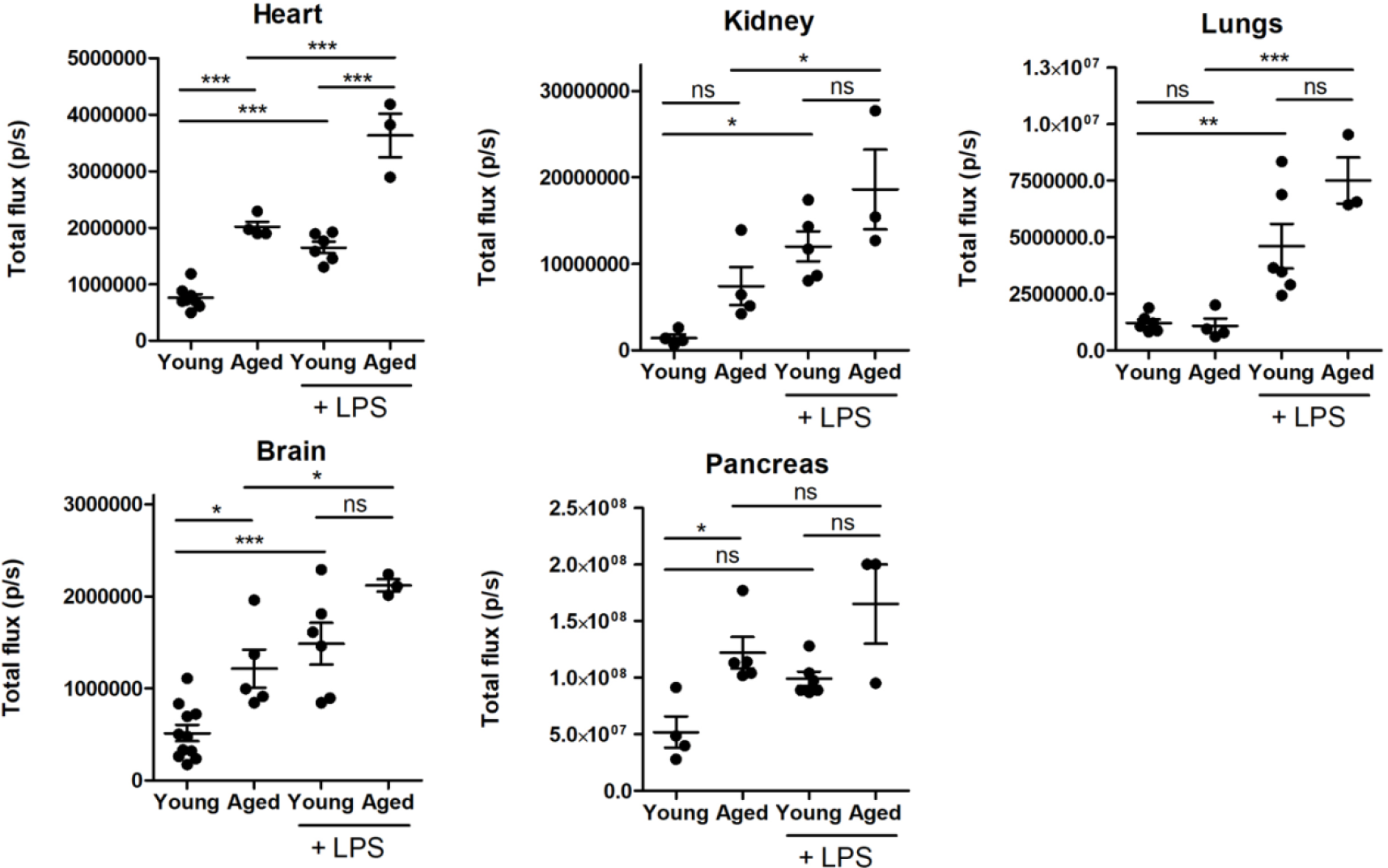
Endotoxemia induces increases in inflammasome activation in young and old tissues: Tissues were extracted from young and aged mice and tissue bioluminescence was measured *ex vivo* (C-D). * = p<0.05, ** = p<0.005, *** = p<0.0005 One-way ANOVA and Bonferroni post hoc test.

To further characterize age-dependent changes in the brain, we utilized caspase-1 reporter mice with cranial windows to visualize inflammasome activation in the CNS *in vivo*. We have recently used this approach to monitor inflammasome activation in the CNS following mild traumatic brain injury (38). Skulls of young and aged mice were thinned completely, and a layer of transparent glue was applied. Windows remain optically clear for up to a year, allowing the continued monitoring of luciferase activity in the brain of living mice. We injected young and aged mice with LPS and monitored CNS inflammasome activation *in vivo*. As expected, we measured increased biosensor activation in young and aged mice treated with LPS, relative to the untreated controls (**Fig. 5A**). However, since both the cranial window surgery and LPS injection increase mortality of aged mice, we also monitored biosensor activation in organotypic brain slices *ex vivo*. We observed increased biosensor activation in slices derived from aged mice that was exacerbated following LPS exposure (**Fig. 5 C-D**). Surprisingly, we did not observe a significant increase in biosensor activation in CNS slices from young mice following LPS addition, suggesting that peripheral chemokines or other inflammatory mediators drive this response following LPS injection *in vivo*. Together, these data indicate that endotoxin challenge in aged mice results in amplified inflammasome activation in the brain.

**Figure. 5.**
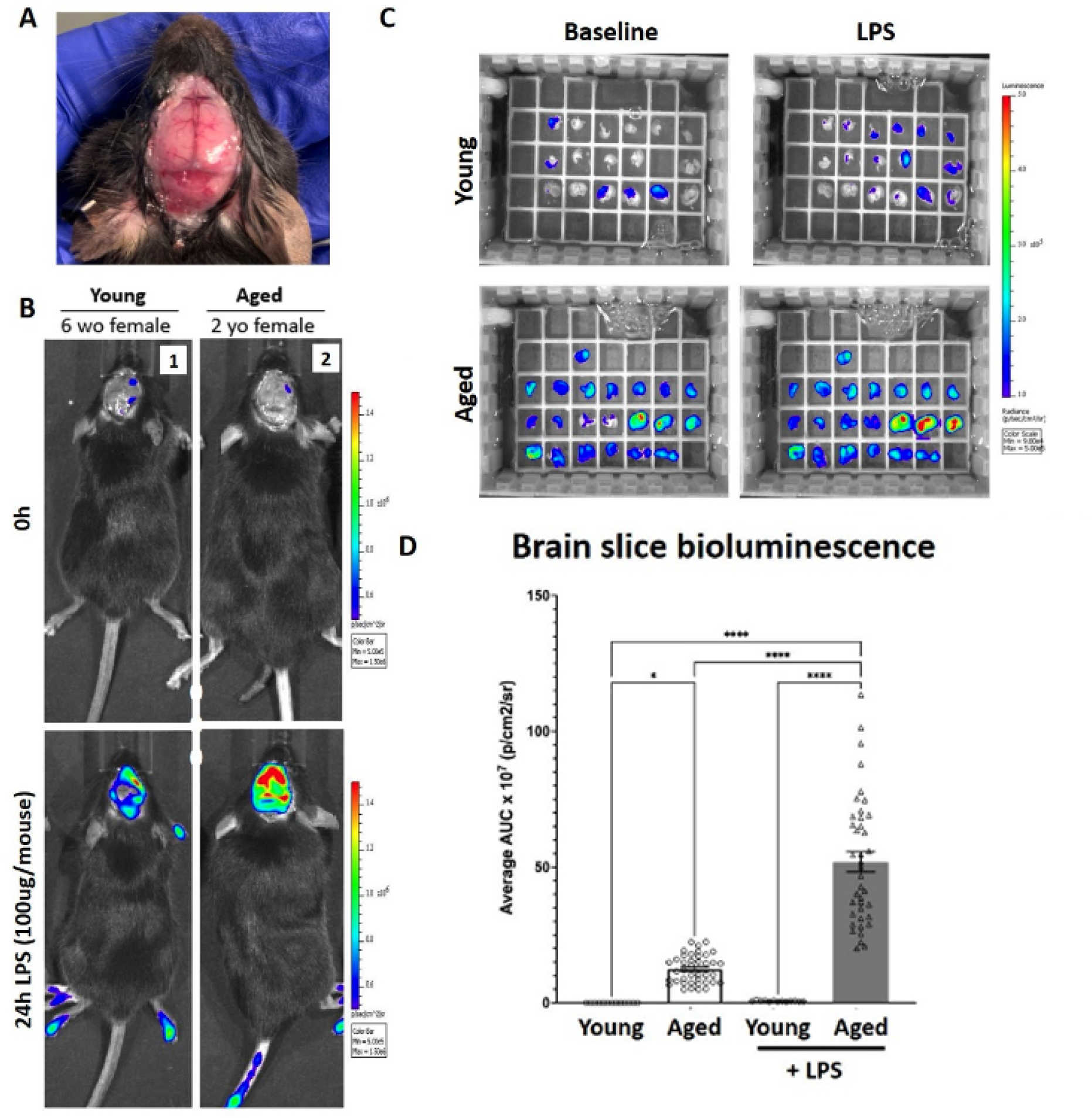
Aged mice exhibit increased inflammasome activation in the brain in response to endotoxin challenge. Representative image of a mouse 6 months following skull-thinning cranial window surgery (A). Young (6-12 week old) and aged (18 month – 2 year old) mice were challenged with LPS (100 μg, i.p). Representative *in vivo* IVIS images prior to (0h) and 24h-post LPS administration (B). (C). Brains were excised from young (3-5 month old) and aged (22 – 23.5 month old) caspase-1 biosensor mice and 400-450μm coronal sections were generated (∼25-45 slices per group). Slices were treated with and without LPS and imaged using the IVIS (D). (C-D) N = 6-10 animals per group, * = p<0.05, *** = p<0.0001 One Way ANOVA, Tukey’s HSD (F-G).

## Discussion

While most work to date has focused on systemic changes in inflammasome signaling during aging, altered inflammatory pathways in specific organs relevant to an age-dependent disease, or aberrant inflammatory pathways activated in specific immune cell populations, the objective of this work was to broadly assess age-associated changes in inflammasome activation in major organ systems and compare this to changes in inflammatory transcripts occurring in these same tissues. To do this, we utilized transgenic mice expressing a bioluminescent reporter of caspase-1, which allowed us to determine which organs and tissues exhibit the strongest degree of age-associated inflammasome activation. We observed elevated caspase-1 activation *in vivo* and in extracted organs *ex vivo*, including the pancreas, heart, kidneys, and brain (**Fig. 1-2, Supplement 1**).

Once we identified tissues with age-dependent increased caspase-1 activity, we further interrogated alterations in inflammatory transcriptional signatures in these tissues. A number of prior studies have performed single-cell or bulk RNA sequencing on aging tissues and demonstrated upregulation of innate immune and inflammatory signaling pathways in aged tissues (18, 49-51). However, when we compared changes in inflammatory transcripts, including transcripts commonly measured to monitor inflammasome upregulation, we observed that these changes did not correlate to the degree of biosensor activation observed in these organs. For example, we observed the most consistent and statistically significant increase in inflammasome associated transcripts in the lungs (**Fig 3)**, despite not observing increased biosensor activation in the lungs (**Fig 1D**). However, we did measure a strong increase in biosensor activation in the lungs of both old and young mice following LPS injection (**Fig 4**), demonstrating that substantial inflammasome activation can occur in the lungs and that this response can be reliably measured using this model. We measured an increased trend in the upregulation of *NLRP3* and *pro-IL-1β* transcripts that was not significant for the number of mice used in these studies, whereas procaspase-1 transcripts were significantly upregulated in most aged tissues (**Fig 3**). Our results demonstrate that age-dependent increases in caspase-1 activation in the heart, brain, and kidneys occurred in the absence of large/significant transcriptional changes in inflammatory pathways in these tissues. An important caveat to our approach is that it relied on bulk RNA isolation, which could obscure cell-type specific changes in inflammatory transcript expression.

We wanted to further assess age-associated changes in inflammasome/caspase-1 activation in response to an inflammatory stimulus. We hypothesized that caspase-1 activation would be exacerbated in response to endotoxemia in aged mice. Despite a relatively small number of animals available for these experiments, we observed statistically significant increases in biosensor activation in the heart, kidney, lungs and brain (**Fig 4**). We also attempted to monitor CNS inflammasome activation directly using cranial windows. These data corroborated our *ex vivo* findings (**Fig 5B**), although a small number of aged animals surviving the window surgery and LPS injection precluded assessment of statistical significance. We also observed a significantly increased response to LPS in brain slice cultures from aged mice (**Fig 5C-D**). Notably, we did not observe an increase in biosensor activation in LPS-treated brain slices from young animals, compared to pretreatment levels (**Fig 5 C-D)**. Since we observed a significant increase in young brains imaged *ex vivo* following LPS injections (**Fig 4**) and observed a similar pattern in young mice with cranial windows (**Fig 5B**), this suggests that peripheral inflammation is required to drive CNS inflammation in young mice. In this context, future studies are required to understand the differential responsiveness of aged CNS tissue to LPS, which could be due to an increased number of peripheral immune cells present in the CNS of aged animals or differential responsiveness of CNS resident cells.

In these studies, we also tried to validate caspase-1 activation in tissues using western blot to detect cleaved caspase-1. However, the presence of age-associated immunoglobulins precluded the detection of cleaved caspase-1. Broadly speaking, the reliable detection of cleaved caspase-1 has been limited to two commercially available antibodies to caspase-1. The first, a rabbit polyclonal antibody, is no longer commercially available, leaving most studies reliant on a commercially available mouse monoclonal antibody to caspase-1. However, as noted in another study (52), the similar size of murine IgG immunoglobulin components to immature and mature caspase-1 confounds detection of caspase-1 in some experimental model systems. We find that aging studies are particularly impacted by this technical limitation due to the comparatively high amount of immunoglobulin deposition observed in aged tissue **(Fig S1**). Although we were able to validate the biosensor signal observed in aged animals using the NLRP3 inhibitor MCC950 (**Fig 2**), the prevalence of age-associated immunoglobulins in the organs examined demonstrates that detection of caspase-1 cleavage using this antibody should be avoided and that secondary antibody controls are required in studies of inflammasome activation in aged animals.

Although our studies and other studies clearly demonstrate a role for the,NLRP3 inflammasome in age-associated inflammasome activation, these studies to not exclude the role of other inflammasomes in this response. Most inflammasomes promote the cleavage of gasdermin D or gasdermin E, which leads to the formation of pores on the plasma membrane that are able to drive changes in cytoplasmic ion concentrations, including K+ efflux, which is known to promote NLRP3 activation (32). Thus, although we observe that a substantial amount of inflammasome activation occurring in aged animals is sensitive to NLRP3 inhibition, it is possible that other inflammasomes initiate inflammasome activation in some organs or tissues, and that these responses induce subsequent activation of the NLRP3 inflammasome.

## Conclusion

In this study, we demonstrate that transgenic mice expressing a caspase-1 biosensor are an attractive model to study age-associated inflammasome activation in mice. In a relatively small cohort of animals, we were able to detect statistically significant increases in inflammasome activation in many organs. Moreover, we observe that inflammasome activation in many organs is not correlated with transcriptional changes of inflammasome associated transcripts, demonstrating the importance of measuring activation of this post-translational inflammatory pathway directly or indirectly, as we have done here.

## Materials and Methods

### Mice

Caspase-1 biosensor mice and C57Bl/6J (strain #000664, Jackson Laboratory) were used for these studies. Mice were bred in house and maintained in pathogen-free conditions at Loyola University Chicago. All experiments were performed in accordance with protocols approved by Loyola University Chicago’s Institutional Animal Care and Use Committee.

### IVIS measurements

For IVIS imaging, caspase-1 biosensor mice were weighed and injected intraperitoneally with 150 mg/kg VivoGlo Luciferin (Promega). Mice were anesthetized with 2% isoflurane/air mixture and imaged 10 minutes following administration of the luciferase substrate using the IVIS 100 Imaging system (Xenogen). For *ex vivo* imaging, tissues were extracted from sacrificed mice, placed in a solution of diluted luciferase substrate (300 μg/mL) and imaged. Bioluminescent images were acquired and analyzed using Living Image software (PerkinElmer).

### Western blot

Tissues isolated from young and aged caspase-1 biosensor mice or C57Bl/6J mice were flash frozen. Tissues were homogenized using an electric homogenizer in lysis buffer (1% NP-40, 100 mM Tris, pH 8.0, and 150 mM NaCl) containing a protease inhibitor mixture (Sigma Aldrich) and shaken on ice for 30 mins. Lysates were collected following centrifugation and protein content was quantified by BCA (Pierce; Thermo Fisher Scientific). Samples were mixed with 2x Laemmli sample buffer and boiled for 5 mins at 95°C. Equal amounts of protein were loaded into a 4-15% gradient gel (Bio-Rad) and transferred onto a nitrocellulose membrane (Bio-Rad), which were blocked with 5% milk for 1 hour and subsequently incubated with anti-mouse caspase-1 (Adipogen) or anti-β-Actin (Santa-Cruz Biotechnology) antibodies. Chemiluminescence was measured using SuperSignal West Femto Chemiluminescent Substrate (Thermo Fisher Scientific) and a FluorChem E machine (Protein Simple).

### RNA isolation and qRT-PCR

Tissues were homogenized by electric homogenization in TRIzol™ (Invitrogen) and RNA was extracted following the manufacturer’s instructions. cDNA was synthesized using the GoScript Reverse Transcription System (Promega) and quantitative real-time PCR was conducted as described previously (40).

### Cranial window skull thinning surgery for *in vivo* CNS imaging

Mice were anesthesized with Avertin and fixed in a small animal stereotaxic system under a dissecting microscope. The skin and periosteum layers were excised to expose the skull, and the skull was thoroughly cleaned. Skulls were thinned using a Microtorque Foredom K. 1070 drill (Foredom Inc.). The drill bit was angled parallel to the skull surface and the microdrill was moved uniformly from anterior to posterior. Every few minutes, the skull was irrigated with saline and dried with compressed air and cotton swabs and the drilling motion was repeated until the skull was optimally thin and the microvasculature was clearly visible. A thin layer of cyanoacrylate glue (C1000, Ted Pella Science) was applied to provide protection, followed immediately by a drop of insta-set accelerator (Bob Smith Industries Incorporated) so that the glue would dry transparent. Animals were allowed to recover for two weeks prior to baseline IVIS imaging and LPS exposure.

### Statistical analysis

Statistical significance was determined using a student’s t test when comparing two groups or one-way ANOVA followed by a Bonferroni multiple comparisons test when comparing multiple groups using GraphPad Prism software (GraphPad Software, Inc.).

### Brain slice preparation and imaging

Mice were anesthetized with 125–250 mg/kg tribromoethanol (intraperitoneally) and decapitated with scissors. Whole brain was then removed and immediately placed in an ice-cold 4 °C oxygenated sucrose artificial cerebral spinal fluid (s-ACSF) solution (206 mM/L sucrose, 2 mM/L KCl, 1 mM/L MgCl_2_, 2 mM/L MgSO_4_, 1.25 mM/L NaH_2_PO_4_, 26 mM/L NaHCO_3_, 10 mM/L D-glucose, 1 mM/L CaCl_2_). Brains were placed in 4 °C s-ACSF and sectioned with a vibrotome at 350–400 μm thick (Leica VT1200S; Leica, Nusslock, Germany). Collected slices were then incubated at 32 °C for 1 h in i-ACSF (124 mM/L NaCl, 3 mM/L KCl, 2 mM/L MgSO_4_, 1.25 mM/L NaH_2_PO_4_, 26 mM/L NaHCO_3_, 10 mM/L d-glucose, 1 mM/L CaCl_2_) before imaging sessions. Each experimental group contained 3–4 animals. Brain slices were imaged at a 2-min interval for a total of 8 min for each series. 100 μL of d-luciferin (20 mg/mL) was added to each well one minute prior to the start of the first imaging series to a final concentration of 2 μg/mL. 100 μL of MCC950 (10 mg/mL) was added to the same chamber up to a final concentration of 1 μg/mL 5 min prior to the second imaging series.

---

**Fig. S1.**
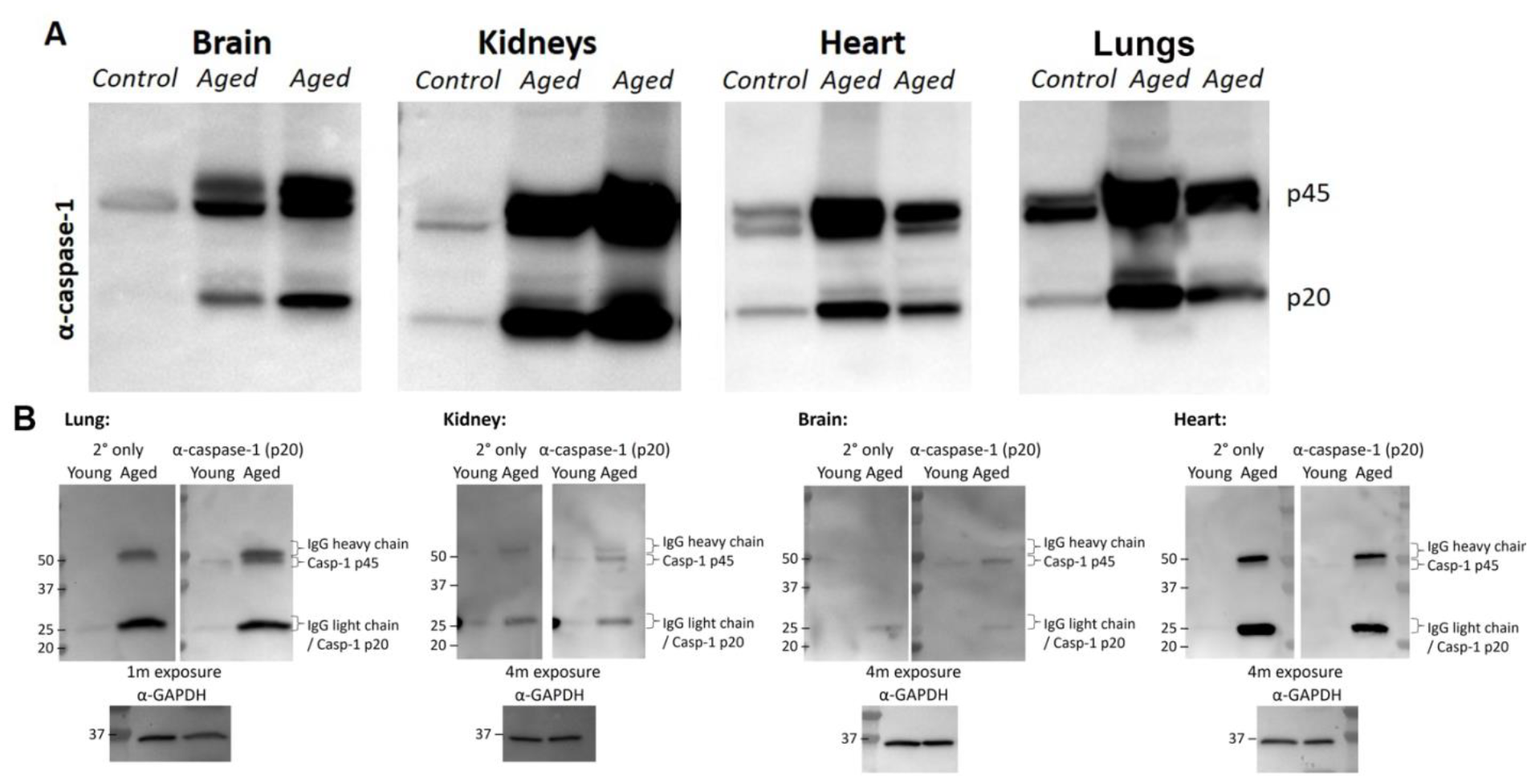
Apparent increases in immature and mature caspase-1 in kidneys, heart, and brain tissue in aged mice. (A) Kidneys, heart, and brain tissues were isolated from young (6-12 week old) and aged (18-24 month old) mice. Tissues were homogenized, lysed, and the expression of caspase-1 was measured by western blot. (B) The same tissues were comparatively analyzed via western blot using α-caspase-1 primary antibody or secondary only control.

**Fig. S2.**
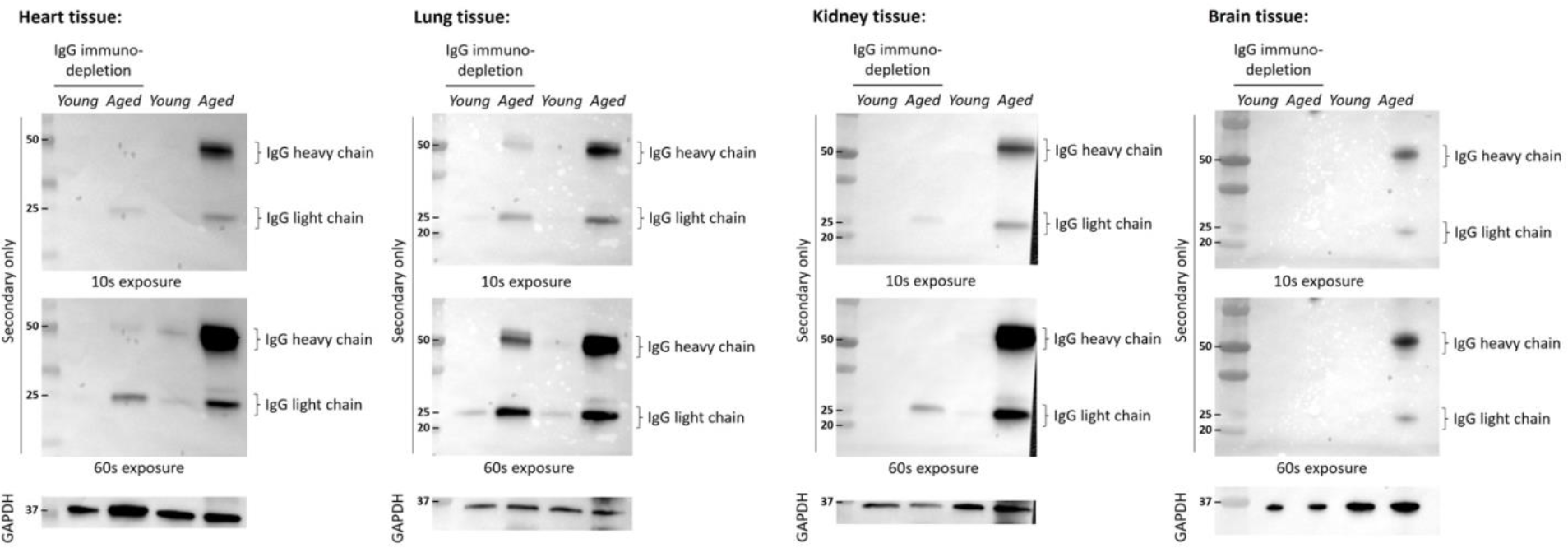
Comparative analysis of IgG immunodepleted tissue lysates with non-immunodepleted tissue lysates. Heart, lung, kidney, and brain tissues of young and aged mice were immunodepleted using agarose Protein A/G magnetic beads (Pierce) and compared to non-depleted lysates from the same mice via western blot utilizing an α-mouse IgG secondary antibody conjugated with HRP (Thermo Fisher Scientific).

**Fig. S3.**
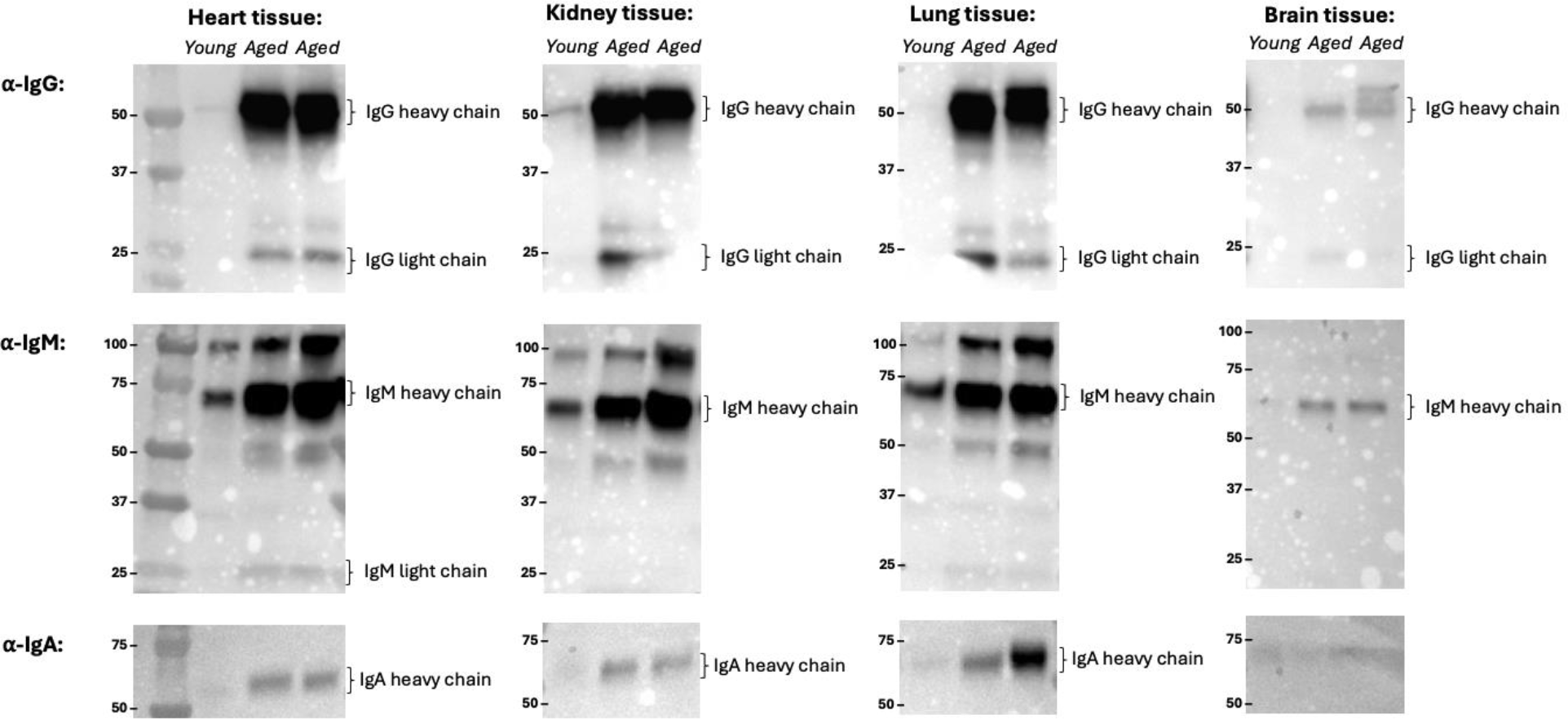
Examination of nonspecific antibody accumulation in aged mouse tissue lysates. Heart, kidney, lung, and brain tissues isolated from young and aged mice. Following homogenization and lysis, the lysates were run via western blot. To analyze the antibody content of the nonspecific antibody accumulation secondary antibodies against IgG, IgM, and IgA were used to block each blot.

**Fig. S4.**
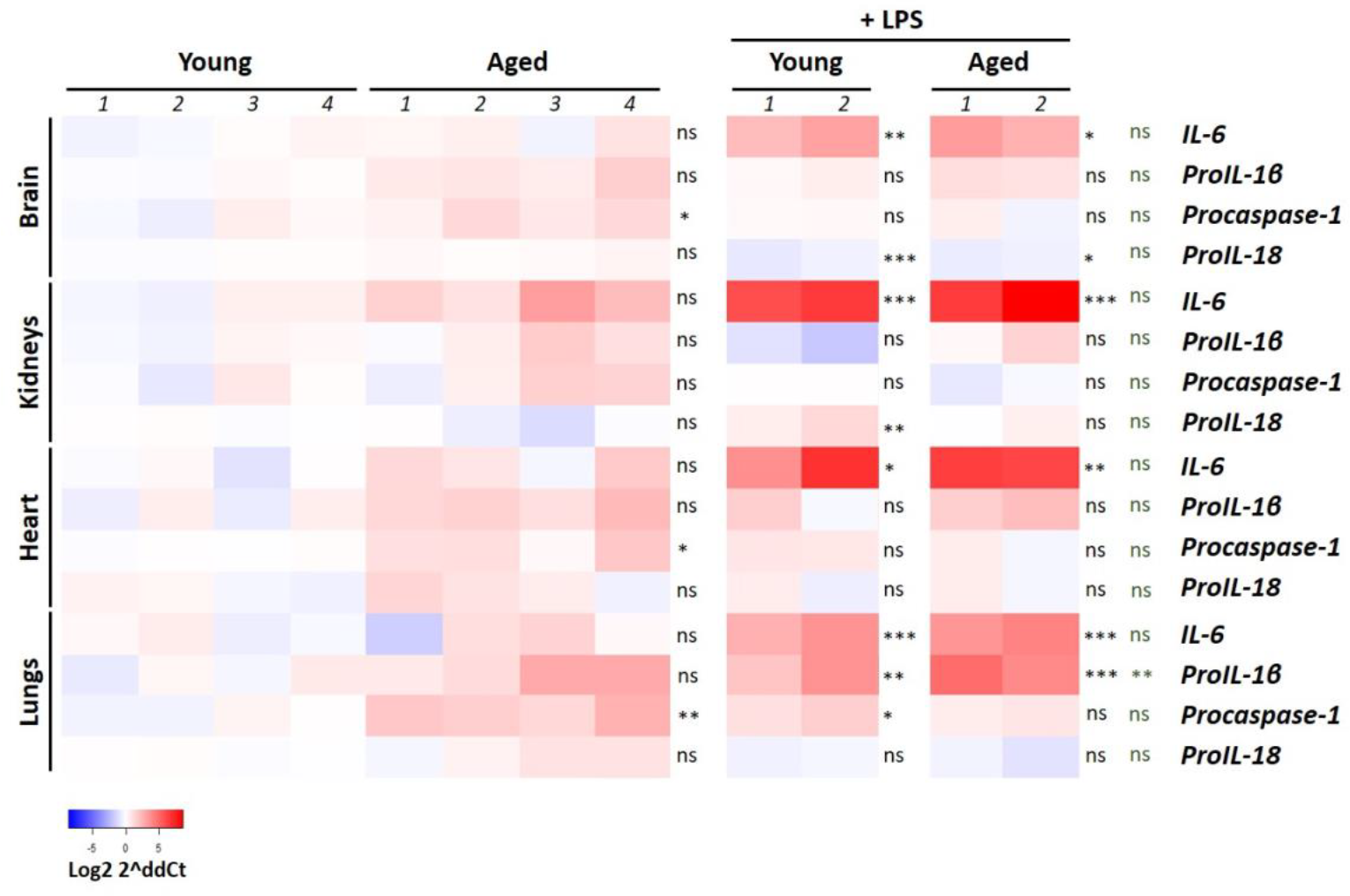
Transcriptional changes in young vs. aged tissues induced by LPS. Heat map depicting expression of the indicated proinflammatory transcripts in tissues isolated from young and aged mice with and without LPS (100 μg, i.p, 24h). Statistics column 1 = young vs. aged, column 2 = young vs. young + LPS, column 3 = young vs. aged + LPS, column 4 = aged vs. aged + LPS. * = p<0.05, ** = p<0.005, *** = p<0.0005 One-way ANOVA and Bonferroni post hoc test.

